# Predicting Adverse Drug Reactions of Two-drug Combinations using Structural and Transcriptomic Drug Representations to Train a Artificial Neural Network

**DOI:** 10.1101/2020.06.30.176016

**Authors:** Susmitha Shankar, Ishita Bhandari, David T. Okou, Gowri Srinivasa, Prashanth Athri

## Abstract

Adverse drug reactions (ADRs) are pharmacological events triggered by drug interactions with various sources of origin including drug-drug interactions. While there are many computational studies that explore models to predict ADRs originating from single drugs, only a few of them explore models that predict ADRs from drug combinations. Further, as far as we know, none of them have developed models using transcriptomic data, specifically the LINCS L1000 drug induced gene expression data to predict ADRs for drug combinations. In this study we use the TWOSIDES database as a source of ADRs originating from two-drug combinations. 34,549 common drug pairs between these two databases were used to train an artificial neural network (ANN), to predict 243 ADRs that were induced by at least 10% of the drug pairs. Our model predicts the occurrence of these ADRs with an average accuracy of 82% across a multi fold cross validation.

Source Code and input dataset used in this study can be found at: https://bitbucket.org/ishita98/prediction-of-adr/src/master/

## 1. Introduction

The most common definition, provided by the WHO, for an Adverse Drug Reaction (ADRs) is “*A response to a drug which is noxious and unintended and which occurs at doses normally used in man for prophylaxis, diagnosis, or therapy of disease, or the modification of physiological function.are noxious and unintended responses to drugs, when they are administered in their normal recommended dosages*”. This study focuses on the prediction of ADRs caused by the drug-drug interaction (DDI) of two-drug combinations. Apart from contributing to various productive drug design strategies like drug repurposing (Zhou et al., 2015), co-administered drugs can exhibit *synergistic DDIs* (Liu et al., 2017), which comprises a new ADR that may be associated with either of the drugs, or the aggravation of an existing ADR. In this study we have proposed an Artificial Neural Network (ANN) that predicts this specific subclass of ADRs using transcriptomic data, compound chemical fingerprint and GO ontologies.

### 1.1 Prevalence and Salient Statistics

Older studies implicate ADRs to be one of the top 10 causes of fatality in the US (Lazarou et al., 1998), while more recent studies report that they account for anywhere for 2.7% to 15% of hospitalizations(Liu et al., 2017; Miguel et al., 2012). Drug-drug interactions account for upto 30% of ADRs (Iyer et al., 2014), and are known to affect clinical outcomes in 80% of those observed in cancer patients (Beijnen & Schellens, 2004). While these statistics are for the US, they are known to be important in all healthcare systems. In France, DDIs are estimated to be the cause of hospitalization in the elderly in as high as 2-5% of the cases, and upto 1% in the general population (Létinier et al., 2019). As a further example, observational studies conducted in a single tertiary care hospital over a period of 6 months reported 598 drug related problems, of which 55.5% were due to drug interactions in 332 patients, of which 39 were putative ADRs (George et al., 2015). Another study focusing on geriatric patients, conducted over a period of one year, observed the incidence of polypharmacy in 22.9% of the patients and 29.5% of the patients were exposed to potentially inappropriate medication (since they were on multiple drugs) (Rajeev et al., 2018). Similarly, studies conducted in several other parts of India have estimated the incidence of suspected ADRs to be nearly 2-3%, among hospitalized patients. Globally, the prevalence of ADRs to the proportion of inpatient hospitalizations were reported to be 2.3%, 4.8% and 7.3% for England, Germany and the United States, respectively (Mulchandani & Kakkar, 2019). The administration of poly-pharma compounds increases the risk of adverse events due to drug-drug interaction (Iyer et al., 2014). If each of the drugs contributes to the cumulative probability of the observed ADR, the overall risk due to multiple drugs increases. To compound this problem, it is believed that 90% of ADRs go unreported (Raj et al., 2019).

### 1.2 Artificial Intelligence in Predicting ADRs caused by DDIs

Computational approaches to predict Adverse Drug Reaction are based on the guilt-by-association rule. The predictions are based on the similarity of molecular structures, molecular circuits and common targets and pathways. In general, these models process multi-dimensional data that are correlated to multi-drug adverse events, and are able to collectively use integrated, heterogeneous databases. These analyses can potentially result in robust and comprehensive models that are precursors to experimental ADR studies that reduce cost and time to perform these studies experimentally, flag interactions that can be fatal and significantly reduce losses from post-marketing surveillance (Chen et al., 2016). Artificial intelligence (AI) algorithms have been used to both detect and classify ADRs for both single and two-drug conditions (see (Lee & Chen, 2020) for an exhaustive review of previous studies for all combinations of the above). Most of these detect and quantify the probability of interaction/ interaction-strength of DDIs, as opposed to a very small number of studies that assign a specific causal relation between the ADR and the DDI in question. We found three studies that use artificial intelligence/ machine learning related algorithms to predict associations of ADRs with DDI pairs. DeepDDI (Ryu et al., 2018) uses a Deep Neural Network (DNN) to predict 86 specific ADRs using structural representation (SMILES) of the drug-pairs using DrugBank’s DDI (Wishart et al., 2018). Decagon (Zitnik et al., 2018) developed a graph convolution neural network that modeled the prediction of polypharmacy as a multi-relational link prediction problem between the drugs, the target and the side effects. They use STITCH (Szklarczyk et al., 2015) as a source for drug-protein interactions, and SIDER (Kuhn et al., 2016) to retrieve side effects of individual drugs, as well as two-drug combinations. Finally, Zheng et al (Zheng et al., 2018) built a machine learning algorithm based on what they term as Highly-Credible Negative Samples. They use drug representation that combines fingerprints, protein targets, pathway enrichment and substituents with ADR labels from the TWOSIDES dataset (Tatonetti et al., 2012). The analysis was designed such that each ADR was predicted individually using binary classification techniques that, historically, were mostly used for DDI detection (Lee & Chen, 2020).

### 1.3 Motivation and Scope

In this study, we present an ANN to predict Adverse Drug Reactions observed when two drugs are co-administered. The independent variable is an encoding of the drug-pair, and the labels for training the ANN are obtained from the TWOSIDES dataset (Tatonetti et al., 2012). The encoded drug data set used, was previously calculated and utilized to predict single-drug ADR (Wang et al., 2016). We have used the same encoding (see Section 2.1), and juxtaposed the signatures of two drugs that constitute the drug-pair whose DDI type we are predicting. In their scheme, which we have also adopted in this study, each drug is represented by its chemical fingerprint, Differential Gene Expression (DEG) signature, and an extension of the response data that represents the enrichment (per drug) for Gene Ontology gene sets. Specific to drug interactions, it should be noted that fingerprints embed structural information in the drug-pair vector and provide an encoding that enables the algorithms to perceive functional group similarities between the two drugs. Nonetheless, using structural characterization of compounds alone can fail in cases where the structural similarities exist, but the side effects differ (Miotto et al., 2018). Wang et al (Wang et al., 2016) have used the LINCS L1000 (Subramanian et al., 2017) perturbagen (including compounds) responses encapsulated by transcriptomic data to predict single drug induced ADRs.

TWOSIDES dataset (Tatonetti et al., 2012) has collated adverse events caused by drug-pairs that can be clearly distinguished from adverse events caused by the individual constituents. The differentiation is either in terms of a completely new type, or a significant enhancement of an existing adverse event when compared to what is caused by a single drug. In effect, in this study and others (Liu et al., 2017; Zheng et al., 2018) the TWOSIDES dataset is used as a source of verified ADRs for combination of two drugs.

While previous studies that predict DDI types (ADRs) from DDI information have proposed that using drug expression data is one of the important and natural next steps (Ryu et al., 2018), there is no study till date that uses it to predict DDI-ADR causal relationships. The use of transcriptomic data supplements the largely dominant target-based method in predicting phenotypic effects. We combine the reported advantages of using multi-label classification to predict multi-drug reactions, in combination with drug induced differential gene expression data (as opposed to only drug-centered features), for prediction of ADRs originating from multi-drug prescriptions.

### 1.4 Artificial Neural Network for Multi-class Predictions

In this study, we use an Artificial Neural Network (Bishop C. 1995) to predict ADRs of two-drug combinations. The Artificial Neural Network (ANN) is an example of supervised learning. In this framework, a classifier attempts to model the relationship between an input and its corresponding output by iteratively minimizing the ‘error’, which is a function of the difference between the predicted output and desired output, over the training set. The layers sandwiched between the input and output layers are called hidden layers. The ANN offers great flexibility through a number of tunable parameters, for example, the number of hidden layers, number of nodes in each layer, error function, optimization method, etc. (Reed & Marks II 1999). This facilitates modeling complex relationships between the input and various output classes. An ANN with ten or more hidden layers is called a deep neural network (DNN) (Goodfellow et al., 2016), with a far greater number of tunable parameters and multiple architectures, making it a powerful classifier. Recent research has demonstrated the efficacy of DNN in predicting single drug ADRs (Uner et al., 2019)., ADRs for drug-drug interactions (Ryu et. al. 2018) and possible ADRs of a new drug (Wang et. al. 2019) among others (Lee & Chen 2020). In this study, we use a lighter-weight ANN with multi-dimensional features that span chemical structure representation, GO ontology terms and drug-induced gene expression (Wang et al., 2016) to predict ADRs induced by two-drug combinations.

## 2. Methods and Materials

The prediction model for ADRs of various drug-pair combinations was structured as a multi-label classification task. The data sources, and the components in the respective resources used in this study are presented in Table S1. The models were built in Intel® Xeon(R) Silver 4114 CPU @ 2.20GHz (40 cores), and 251.5 GiB RAM machine running on Ubuntu 16.04 LTS. The data processing and ANN model creation was done in a Python environment using various libraries, namely, ScikitLearn (v0.22.2.post1) (Pedregosa, G., Varoquaux, A., Gramfort, V., Michel, B., Thirion, O., Grisel, M., Blondel, P., Prettenhofer, R., Weiss, V., Dubourg, et al, 2011), TensorFlow (v2.1.0)(*Tensorflow*) and Keras (v2.3.1) (Chollet, 2015).

### 2.1 Drug-pair Representation and the Associated ADRs

LINCS L1000 data (Subramanian et al., 2017) has differential expression data for a total of 20,338 perturbagens. Zhang et al (Wang et al., 2016) have used this to encode/ represent drugs using 3 components (Table S1). In our study, this representation was directly downloaded from their paper (*Prediction of drug side effects*, 2016.; Wang et al., 2016). Further, the representation for a drug pair was formed by concatenating individual drug encodings. Thus, each input vector has 11,164 (Supplementary Figure 1(b)) features.

Drug-drug-ADR associations to train our models are extracted from TWOSIDES database (Tatonetti et al., 2012). ADRs in the TWOSIDES data are extracted from Adverse Event Reporting System (AERS) from the FDA website.

The categorization of these ADRs use standardized MedDRA terminology. It comprises 868,221 associations between 59,220 pairs of drugs that collectively exhibit an aggregate of 1301 adverse events. The drug names from LINCS L1000 metadata (called *pert_iname*) were used to map the records against the drug names in TWOSIDES. Subsequently, the associated perturbagen ID (*pert_id*) was used to merge the records from the Wang et al dataset. While 206,792,742 drug-pairs had transcriptomic data available from L1000 dataset, only 34,549 pairs had both expression *and* DDI ADR information available from TWOSIDES data (see Figure 1).

**Figure 1:**
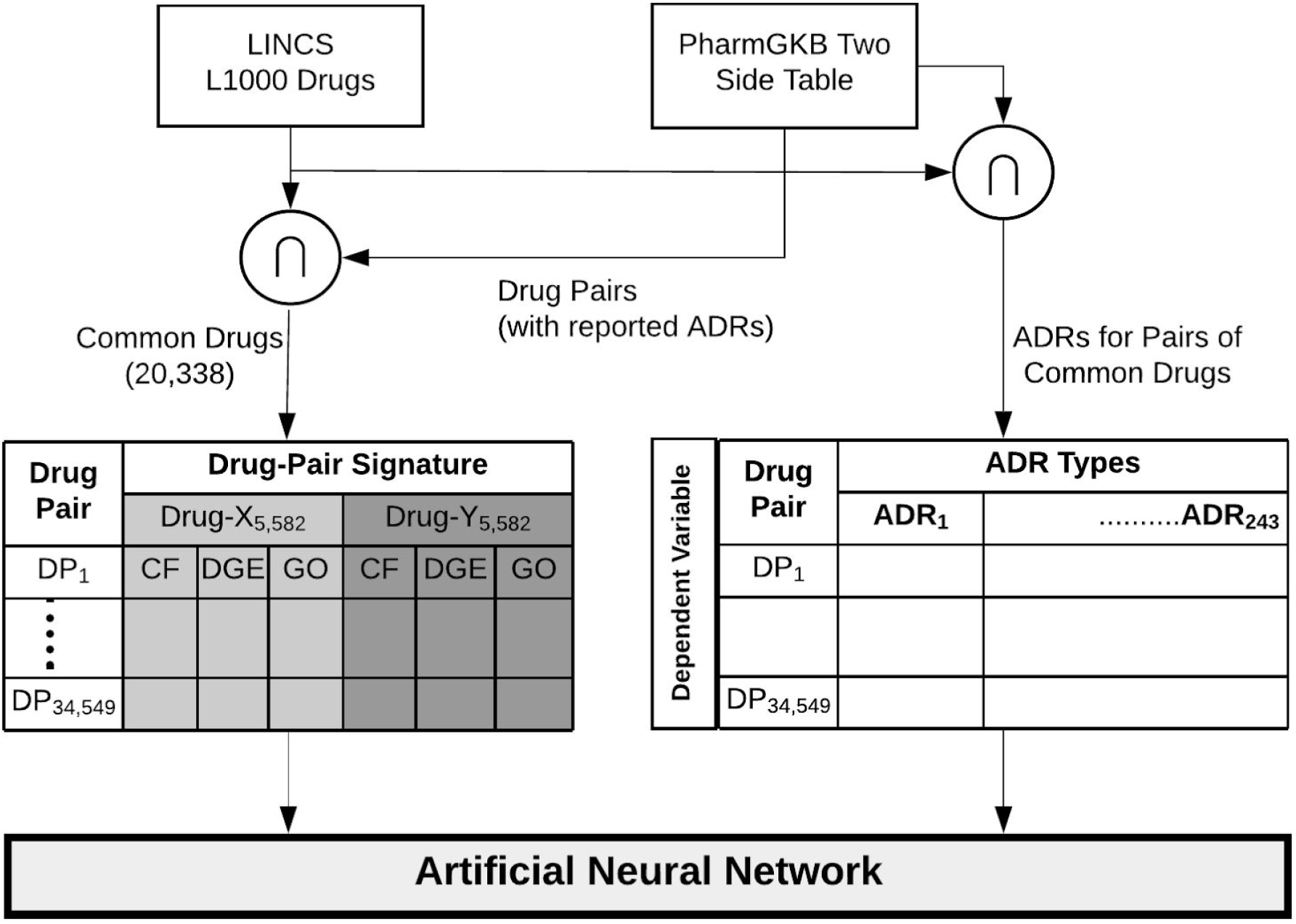
Dataset preparation

Some of the ADRs are under-represented in the dataset, and will tend to skew the model. We decided to have a cut-off based on the minimum number of samples that need to manifest a particular ADR, for the ADR to be considered in the study. We made empirical tests across five different percentages ranging from 10% through 90% at 20% intervals. To elucidate, for the 10% case, at least 10% of the 32,549 input samples is expected to present an ADR for that ADR to be selected in the study. One-hot encoded vectors were used to represent the presence/absence of ADRs collectively associated for a particular drug pair.

### 2.2 ANNs for the Prediction of ADRs for drug-pair combinations

Several machine learning models were tested, and a summary of methods and results is presented in the Supporting Information (Section S3). ANNs were chosen based on performance evaluation. In this model, prediction of ADRs for two-drug combinations has been posited as a multi-label classification problem, since each drug pair can be associated with multiple ADRs. Recent literature has demonstrated the efficacy of artificial neural networks towards predicting ADRs associated with single drug and drug-drug interactions, even with high dimensional and voluminous data encodings (Lee and Chen 2020). The comparatively smaller dataset we use in this study prompted us to use lighter weight ANNs that avoids the challenges associated with high computational complexity. An ANN is a network of nodes or artificial neurons distributed across different layers. A representation of the architecture used in this study is shown in Figure 2. The first layer comprises the input layer, which is the drug pair encodings in this study. In a trained ANN, each input is multiplied by an optimal weight. These weighted inputs are accumulated at each node in the subsequent layer and passed through a nonlinear function, called the activation function. This activation function ensures the weighted linear combination of inputs at any node lies within a specified range such as [0,1], or [−1,1], etc. The output of one layer serves as the input to the next, culminating in the final output layer. Through weights associated with the interconnections of nodes between layers, the network attempts to model the inherently complex boundaries between output classes in the input feature space. The weights in the network are adjusted iteratively in the training phase through a learning mechanism called backpropagation (Bishop C.,1995).

**Figure 2:**
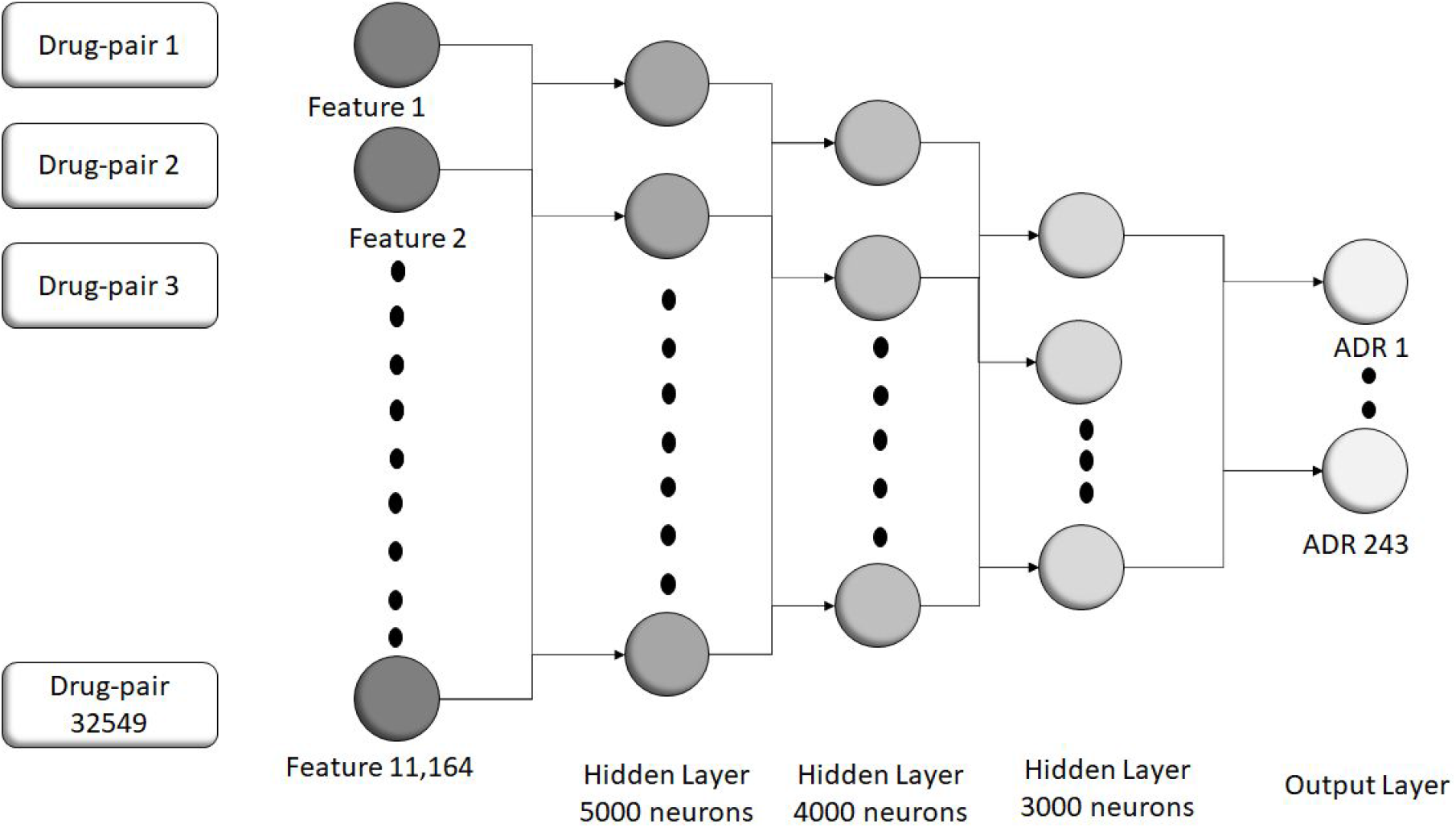
Architecture of the Artificial Neural Network used for the neural model.

#### 2.3.1 Selection of ANN Hyperparameters

The tuning of various parameters is typically empirical (Reed & Marks, 1999). In our model, the ANN has an input layer comprising 11,164 nodes corresponding to the features that represent two-drug combinations, three hidden layers and an output layer that comprises 243 nodes corresponding to the 243 possible ADRs shortlisted for this study (Figure 2). A brief description of the parameters of the ANN and the rationale behind the choice of specific parameter values is presented in Table 3, and complete details of all the experiments that were run to evaluate the optimal values of each of the hyperparameters is explicated in the Section S2.

**Table 3:**
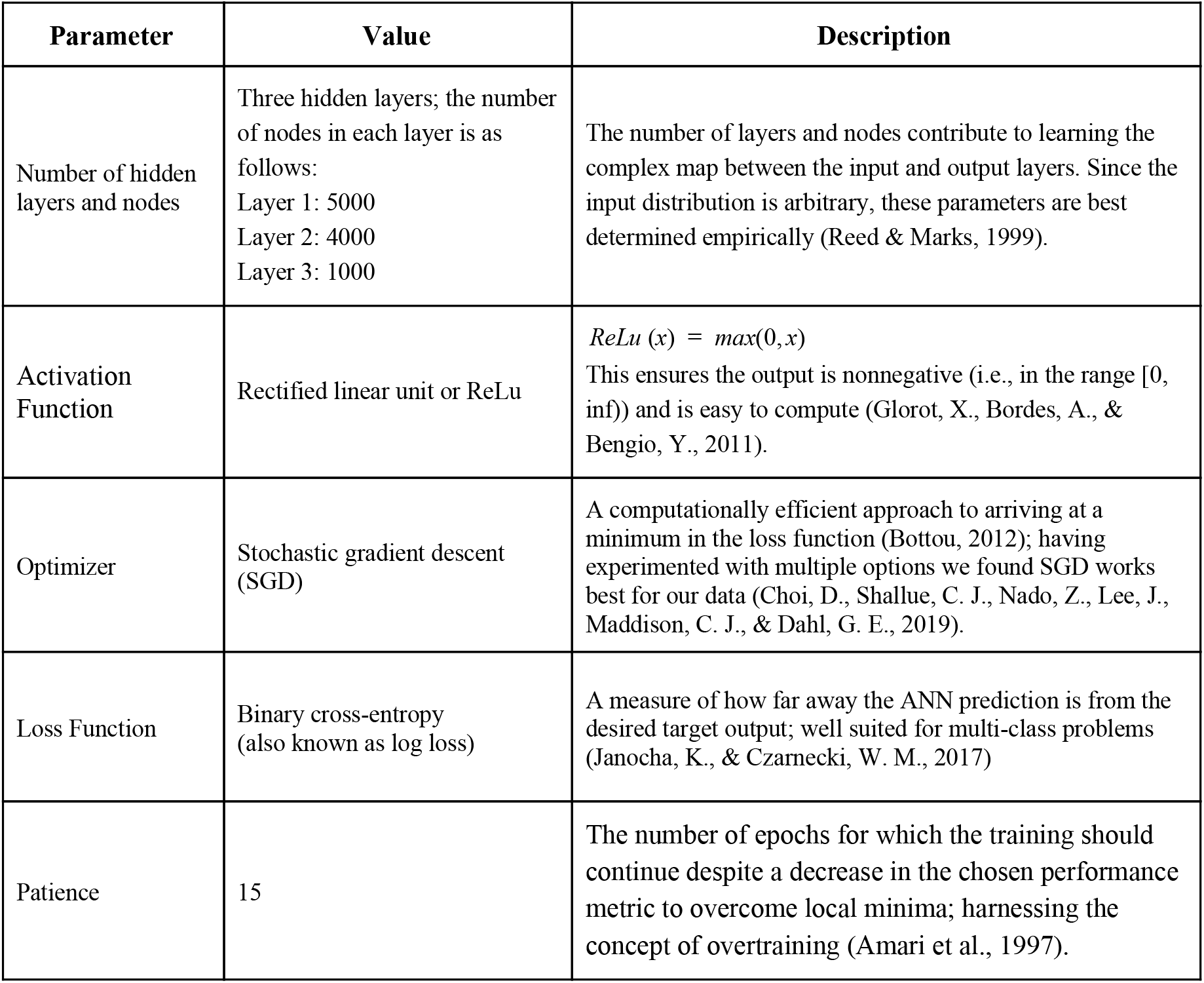
ANN parameters and their values

| Parameter | Value | Description |
| --- | --- | --- |
| Number of hidden layers and nodes | Three hidden layers; the number of nodes in each layer is as follows:<br>Layer 1: 5000<br>Layer 2: 4000<br>Layer 3: 1000 | The number of layers and nodes contribute to learning the complex map between the input and output layers. Since the input distribution is arbitrary, these parameters are best determined empirically (Reed & Marks, 1999). |
| Activation Function | Rectified linear unit or ReLu | $ReLU(x) = \max(0, x)$<br>This ensures the output is nonnegative (i.e., in the range $[0, \infty)$) and is easy to compute (Glorot, X., Bordes, A., & Bengio, Y., 2011). |
| Optimizer | Stochastic gradient descent (SGD) | A computationally efficient approach to arriving at a minimum in the loss function (Bottou, 2012); having experimented with multiple options we found SGD works best for our data (Choi, D., Shallue, C. J., Nado, Z., Lee, J., Maddison, C. J., & Dahl, G. E., 2019). |
| Loss Function | Binary cross-entropy (also known as log loss) | A measure of how far away the ANN prediction is from the desired target output; well suited for multi-class problems (Janocha, K., & Czarnecki, W. M., 2017) |
| Patience | 15 | The number of epochs for which the training should continue despite a decrease in the chosen performance metric to overcome local minima; harnessing the concept of overtraining (Amari et al., 1997). |

#### 2.3.2 Interpreting the ANN Output and Evaluation of the Model

After the training process of the ANN is completed, a validation set comprising data that the network has not seen in the training phase is used to evaluate the model. This process was done in two stages:

a. **Binary Representation**: The last layer of the ANN with the loss function maps the values of each of the 243 nodes in the output layer corresponding to the 243 ADRs to a value in (0,1). This represents the probability that the input that corresponds to a drug pair combination will present that ADR. The target, however, is a binary value that is either a 0 or a 1 that indicates the absence or presence, respectively, of each ADR. Thus, there is a need to map the probabilities at the output layer to binary predictions. This can be done using a ‘threshold’, i.e., a value that sets any value less than itself to 0 and greater than or equal to itself to 1. To find a suitable threshold, we first obtained the model’s prediction for each of the training samples. Then, for each ADR, we generated 100 thresholds between the minimum and maximum probabilities predicted for that ADR. From these 100 thresholds we selected the one that maximized the F1 score for that ADR. These ADR-adaptive thresholds are then applied to the predictions on the validation set to obtain binary predictions. These binary predictions were compared to the ground truth to evaluate the model’s efficacy.
b. **Performance Evaluation and Prediction of ADR for unknown combination**: The model’s efficacy is determined using cross-validation on four separate subsets of the data as follows: (i) From the original set of 32,549 drug pairs, 8,000 drug pairs were randomly sampled and split to form four distinct subsets of 2000 drug pairs each (say - S_1_, S_2_, S_3_ and S_4_) to be used for validation. (ii) For each validation set S_i_, the corresponding training set was created by combining the 24,549 drug-pairs that remain in the main dataset (Ds), with 3 subsets S_j_ (with j≠i) using the set S_i_ with unknown drug pair combinations as the validation set. (iii) For each training set, the performance was evaluated on the corresponding validation set. The final performance is reported as an average of the performances on the 4 validation sets. The macro-averaging values of accuracy, RoC and F1-score are used as evaluation metrics as implemented in the Keras API (look in the Supporting Information of (Ryu et al., 2018) to find the equations for the above).

## 3. Results

### 3.1 Comparison Between the Five Input Datasets Constructed Based on Number of Samples Manifesting a Particular ADR

Empirical studies to decide on the sample cut-off percentages that manifest ADRs show that using a 10% threshold resulted in optimal model prediction (see Section 2.1). 243 ADRs were present with a 10% cut-off, 165 for 30%, 68 ADRs for 50%, 18 for 17% and 3 for 90% (see Figure S2(j) that compares average scores across four cross validated sets for each of the five partitions of data, Figure S2(k) that confirms the 10% partition yields the most number of ADRs, when a 0.65 cut-off is used for the F1-score). These results are consistent with the expectation that the power of discrimination reduces between the data samples with increase in the percentage of drug-pairs that exhibit a particular ADR for it to be selected in the training set.

For 20,338 drugs in the L1000 dataset, 206,827,791 unique pairs were created, of which 34,549 drug-pairs were common to both L1000 and TWOSIDES dataset. As detailed in Section 2.3.2, subsets of 32,549 were used to build the models and the remaining 2000 combinations were used to predict the unknown ADRs. This was repeated for four such training and validation subsets, ensuring that each subset of 2000 combinations used to test the model were randomly sampled and not overlapping with the corresponding training set. Of the 243 ADRs, *Respiratory Distress Syndrome* had the least drug-drug associations (10% association), and *Decreased Arterial Pressure* had the most number of drug-drug pairs associated with it (41% associations). The number of ADRs associated with a drug pair ranged from 0 to 195 out of 243 ADRs, and had an average of 43.19.

### 3.2 Performance of the ANN Model

The architecture of the final version of the ANN is presented in Table 3, and a detailed analysis of how we arrived at this configuration through empirical testing is in Section S2. The ANN yielded an average accuracy of 82% across all 243 ADRs. The top ADRs predictions had a minimum RoC of 0.68, after which the values dropped by larger quantum. These top ADRs are presented in Table 4. The boxplot for the measured macro-evaluation metrics for all ADRs is presented in Figure S4 (Supporting Information). The performance of each of the ADR in terms of the evaluation metrics is provided as Supporting Information (file name: *Evaluation Metrics for the ANN Model.csv*). The maximum and minimum of each of the evaluation metrics, corresponding to the most and least predictable ADRs for each of the performance measures, viz. accuracy, F1-score and RoC, is reported in Table S2.

**Table 4:** Top-performing ADRs

| ADR | Accuracy | Macro F1 Score | Macro ROC |
| --- | --- | --- | --- |
| Drug addiction | 0.864750 | 0.703959 | 0.702692 |
| Bruxism | 0.825875 | 0.702245 | 0.709496 |
| Mod | 0.813875 | 0.692815 | 0.698257 |
| Bursitis | 0.885625 | 0.691321 | 0.699877 |
| Fibromyalgia | 0.859375 | 0.690880 | 0.693449 |
| Migraine | 0.835000 | 0.689689 | 0.697851 |
| Arteriosclerotic Heart Disease | 0.781500 | 0.687058 | 0.689559 |
| Acid Reflux | 0.776000 | 0.686985 | 0.690390 |
| Elevated Cholesterol | 0.773250 | 0.685067 | 0.687172 |

### 3.3 ADR Predictions

A set of 243 ADRs was identified, where each of those were manifest in at least 10% of the total samples. An F1-score greater than 0.5 ensures positive discrimination of the model. With a threshold of 0.68 on the RoC, we obtain 9 most predictable ADRs out of the 243 ADRs. The macro-averaged evaluation metrics after validating with 4 different datasets for the top-performing ADR is shown in Table 4 (for the complete list of ADRs and performance metrics refer to the file named *Evaluation Metrics for the ANN Model.csv* in the Supporting Information) .

### 3.4 Use Case of Naproxen and Goserelin

An extensive analysis of ADRs observed for multiple drug pairs considered is non feasible, and beyond the scope of this work. Instead we provide the complete list of results as a table in the Supporting Information (file name: *Drug_pair_output_all_subsets.xlsx*). Domain experts can use this to investigate the drug or drug-pair of interest. As with any predictive machine learning/ data mining results, it is unreasonable to expect every ADR identified by the ANN to be significant (Yi Zheng et al., 2016). As an exemplar of the type of analysis that maybe helpful to researchers we highlight the case of ADRs predicted for the Naproxen-Goserelin case since it achieved the highest accuracy score of 92%. As is the case with the results provided as an Excel document in the Supporting Information, each pair of the output data we have reported is from the test set from the respective partition of the data. While the performance metrics indicate the confidence in prediction for each drug-pair, these tables add value to a domain experts’ analysis by indicating other plausible ADRs, not reported by training dataset (TWOSIDES), albeit subject to validation experiments. Naproxen is a nonsteroidal anti-inflammatory drug. It is used to treat acute pain as well as pain related to rheumatic diseases. Goserelin is a synthetic hormone which stops the production of the testosterone in men and may stimulate the growth of cancer cells. The pharmacokinetics, as reported by DrugBank, indicates that Naproxen may decrease the excretion rate of Goserelin, which could, in turn, result in a higher serum levels (Wishart et al., 2018).

For this drug combination, a total of 16 ADRs were predicted to occur. 10 out of 16 ADRs were also reported by the Tattonetti et al dataset. All 6 of the other ADRs were reported by other databases as adverse events for at least one of the drugs in the combination. The resources used to confirm these observations are Drugs.com (*Goserelin Side Effects: Common, Severe, Long Term - Drugs.com*, 2019; *Naproxen Uses, Dosage, Side Effects & Warnings - Drugs.com*, 2020) and SIDER (Kuhn et al., 2016). *Pain* is reported to be caused by both drugs independently, but *neuropathy* was not reported anywhere in the literature. Figure 5 captures the detailed annotation for this drug combination. Both the figure and this format of presenting results were used in earlier studies (for example (Zheng et al., 2018)).

**Figure 5:**
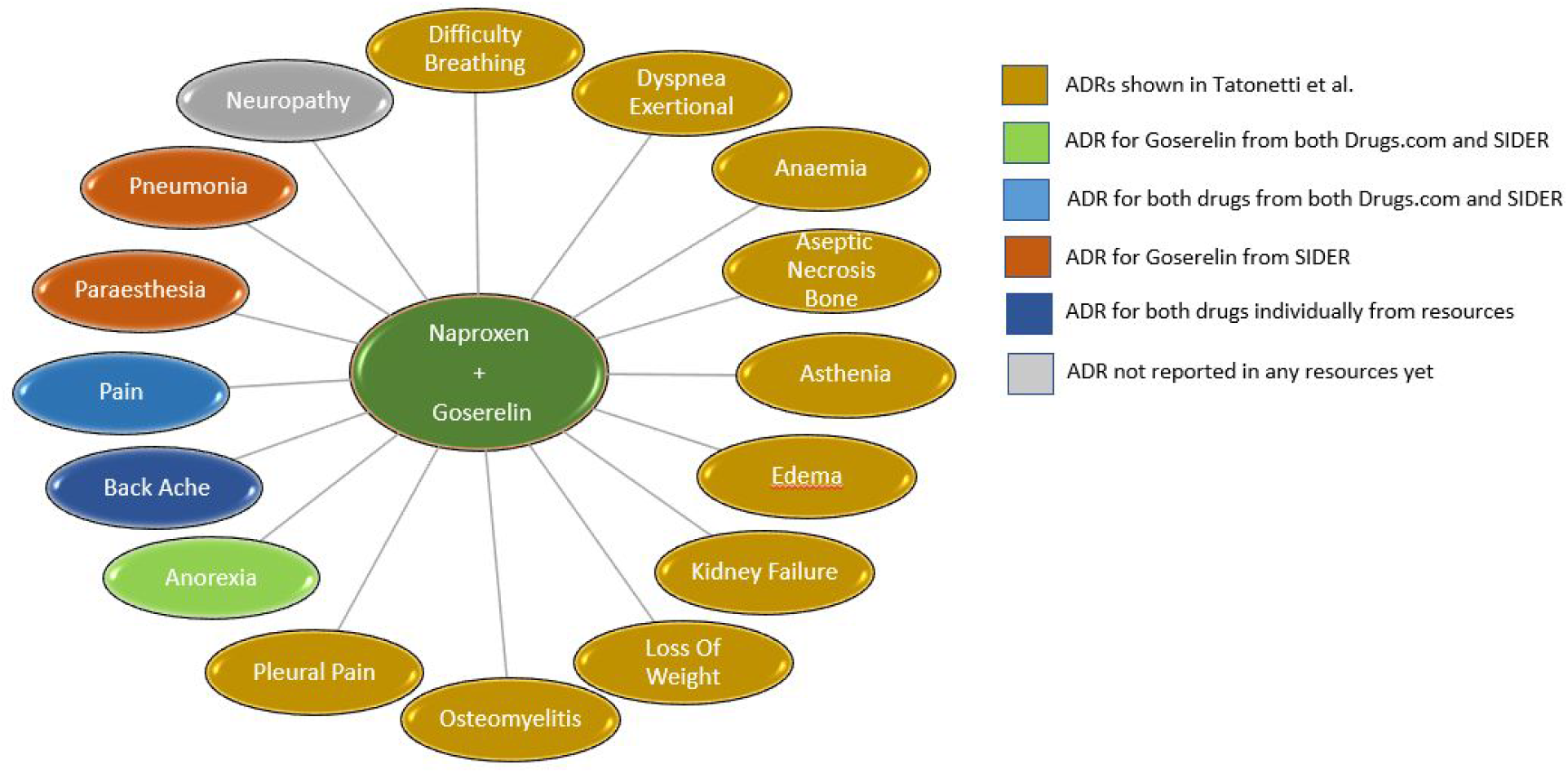
Adverse Drug Reactions predicted by ANN for the combination of Naproxen and Goserelin

## 4. Discussion

In this study, we presented a scalable ANN model to predict Adverse Drug Reaction using transcriptomic data. Even if the transcriptomic signatures are measured individually to any drugs of interest, our model provides ADR estimates to use it in combination with any other drug whose gene expression is available. The use of transcriptomic data reduces the dependency on prior knowledge of protein and drugs targets, which in most cases are incomplete. To our knowledge, drug induced gene expression data has not been used for predicting DDIs, though it has been brought up as a promising method to complement existing predictive studies (Ryu et al., 2018). Transcriptomic data hold direct observation of phenotypic effects of drugs, and is comprehensive as compared to the dominant target-based data used to study ADRs (Wang et al., 2016). As an additional analysis, it would be interesting to compare the results from previous studies for common drug-pairs in future work.

To ensure reproducibility, we have shared all the code, the original input data used to train the model, and the data partition sets needed to reproduce the performance evaluation metrics reported in this study in the supporting information documents. Considering the availability of various high performance python libraries, this study can be easily extended to multiple deep learning and AI algorithms with the provided data sets.

Integrating datasets, even for common biological or chemical entities across highly curated datasets is an intensive, time-consuming exercise (Murali et al., 2020). The scope of this project only covers the inclusion of ADRs resulting from DDIs reported in the TWOSIDES data table. Further improvements through enrichment of observed ADRs for drug pairs can be achieved by integrating the above with DrugBank’s DDI (Wishart et al., 2018), all drug-drug side effects reported in SIDER (Kuhn et al., 2016) and PAERS (Poleksic & Xie, 2019).

## Supporting information

ADR Prediction Comparison

Drug Pairs With True and Predicted Labels

