## Supporting Information for "Predicting Adverse Drug Reactions of Two-drug Combinations using Structural and Transcriptomic Drug Representations to Train a Artificial Neural Network"

### Prediction of Adverse Drug Reaction for combined medication using Transcriptomic data

Susmitha Shankar[1], Ishita Bhandari[2], David T. Okou[3], Gowri Srinivasa[2], Prashanth Athri[1,*]
[1]Department of Computer Science and Engineering, Amrita School of Engineering, Bengaluru, Amrita Vishwa Vidhyapeetham, India
[2]PES Center for Pattern Recognition and Department of Computer Science and Engineering, PES University, Bengaluru, India
[3]Division of Gastroenterology, Hepatology and Nutrition, Department of Pediatrics, Emory University School of Medicine, Atlanta, GA

### Supporting Information

#### Index

1. Input Feature Representation……………………………………………………….Figure S1
2. Tuning of ANN parameters………………………………..……................Figure S2
3. Comparison across the various algorithm………………………...…………..Figure S3
4. Evaluation metrics for the model creates using Artificial Neural Network …….……………………………………………….………...Figure S4
5. Comparison of ANN without any specific negative sample study versus state of art from Zheng et al. …………………………....……....Figure S5
6. Minimum and maximum of each of the evaluation metrics out of 243 ADRs……………………………………………...…….Table S1

#### S1. Input Feature Representation

LINCS L1000 (Subramanian et al., 2017) data comprises 20,338 perturbagen. This is used as the primary data source for drug-drug combinations. We used a subset of all possible combinations of these above perturbangen, depending on whether the combination's Adverse Drug Reaction[s] were available in TWOSIDE dataset (Tatonetti et al., 2012). 34,549 drug pairs out of 206 million had ADRs reported in (Tatonetti et al., 2012) ( Figure S1(a)).

These drug pairs were vectorized as shown in Figure S1(b). Each drug in the drug-pair was represented by the combination of its 166-bit MACCS chemical fingerprints, differential gene expression profile for 978 *landmark genes* (see (Subramanian et al., 2017)) and expression for 4438 gene ontology terms (Wang et al., 2016). Thus, each drug is a vector of size 5582 (166 + 978 + 4438). A drug pair is represented as a concatenated vector of features representing individual drugs (5582 + 5582 = 11164 features).

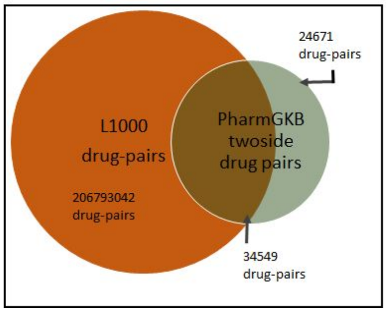


(a)

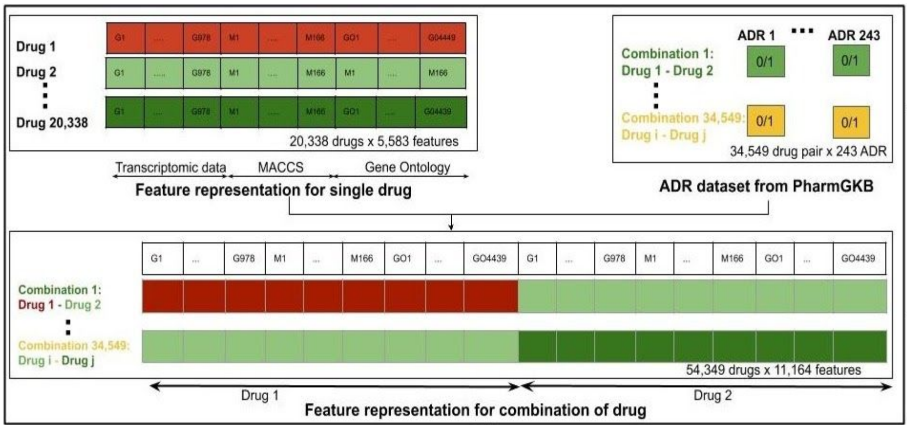


(b)

**Figure S1:** (a) Drug pair origins: A total of 59,220 drug pairs were extracted from PharmGKB dataset for the prediction of Adverse Drug Reactions. Out of these, 58.3% (34549) common drug pairs were used to build the model. (b) Vectorized representation of drug-pairs: A drug is represented by the differential expression, chemical fingerprint and expression level of Gene Ontology terms. The drug pair is represented by the concatenation of each drug vector in the drug-pair.

Table S1: Data sources used in the study

| Source | Description | Download Date |
|---|---|---|
| *Source of:* Drug Encodings<br>*Original Source:* Wang et.al<br>(Wang et al., 2016) | Part:1 Chemical Fingerprint (CF) representation of drugs<br>MACCS fingerprints generated for all drugs<br>No. of Bits: 166<br>URL (for all 3 parts): http://maayanlab.net/SEP-L1000/#download | 01, 2020 |
| | Part 2: Drug response gene expression (DGE)<br>Compound gene expression values from 'landmark genes', as quantified by 'Characteristic Directions' (see original source)<br>No. of genes: 978 | |
| | Part 3: Enrichment p-value per drug calculated for gene sets as created from GO<br>No. of GO terms used: 4,438 | |
| *Source of:* Drug name & ID<br>*Source:* LINCS L1000 data<br>(Subramanian et al., 2017) | Drug name and Drug_ID from LINCS L1000 data<br>These names (pert_iname) and ID (pert_id) were used to map across the Wang et al and TWOSIDES<br>URL: https://www.ncbi.nlm.nih.gov/geo/query/acc.cgi?acc=GSE70138<br>https://www.ncbi.nlm.nih.gov/geo/query/acc.cgi?acc=GSE92742 | 05, 2019 |
| *Source of:* Drug-pair ADRs<br>*Source:* TWOSIDES dataset<br>(Tatonetti et al., 2012) | Adverse Drug Reaction for drug-pairs<br>Number of ADRs: 1,301<br>URL: https://purl.stanford.edu/zq918jm7358 | 01, 2020 |

#### S2. Tuning of ANN Hyperparameters

Tuning of ANN hyperparameters are empirically driven, and we conducted a series of experiments to determine their values based on continuous evaluation of scores (see below) to arrive at perceivably optimal values.

##### S2.1. Number of Layers and Nodes

The number of layers can have a strong bearing on training results. Too many layers can cause overfitting or even cause exploding gradient problems, while selecting a smaller number of layers can cause the model to not train well. Similar is the case with the numbers or nodes in each layer. Arriving at the right combination is key to obtaining high accuracies without overfitting. We report the comparison of three architectures with five, three and two hidden layers. They are referred to as 'High', 'Medium' and 'Low' indicative of the relative computational load in Figure S2(a, b and c).

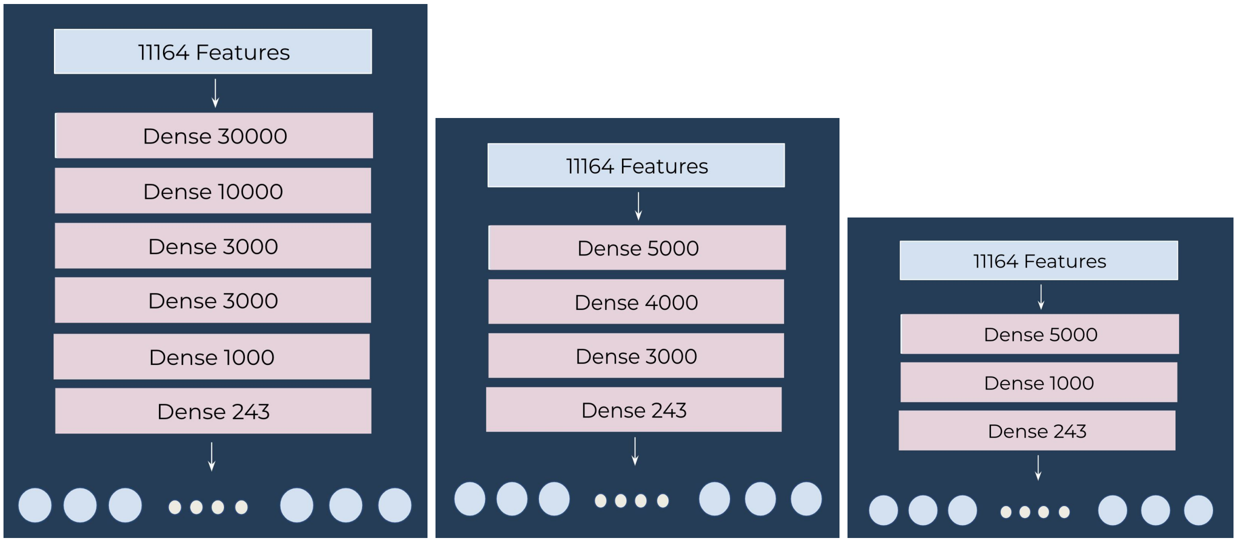


(a) High (b) Medium (c) Low

**Figure S2:** ANN architectures with (a) 5 hidden layers - High (b) 3 hidden layers - Medium (c): 2 hidden layers - Low

.

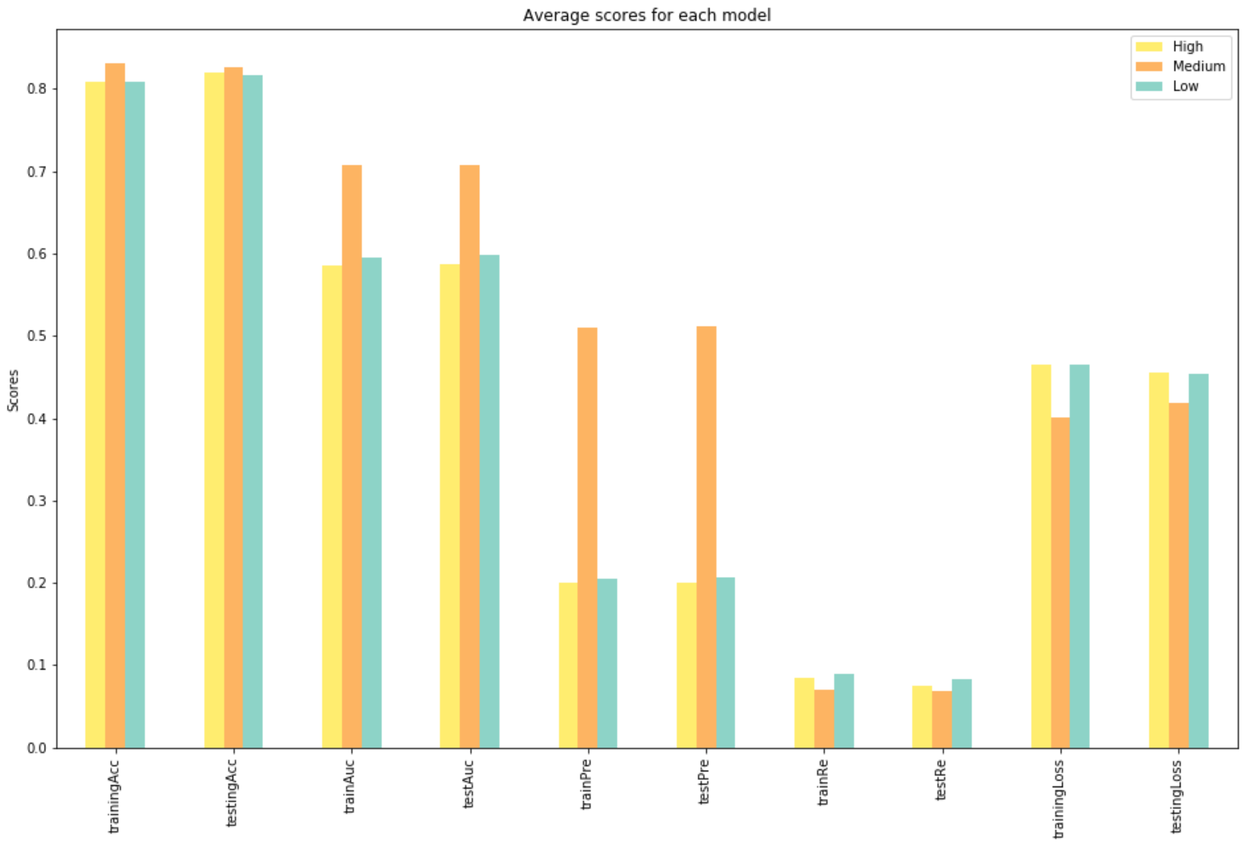


**Figure S2 (d): Performance comparison of the HIgh, Medium and Low models.**

**Figure S2** (d) : results obtained after training of the model using these architectures. It is clear that the "Medium" architecture performs better than the other combinations by almost every performance measure (accuracy, AUC, precision, recall and loss) for training and testing sets.

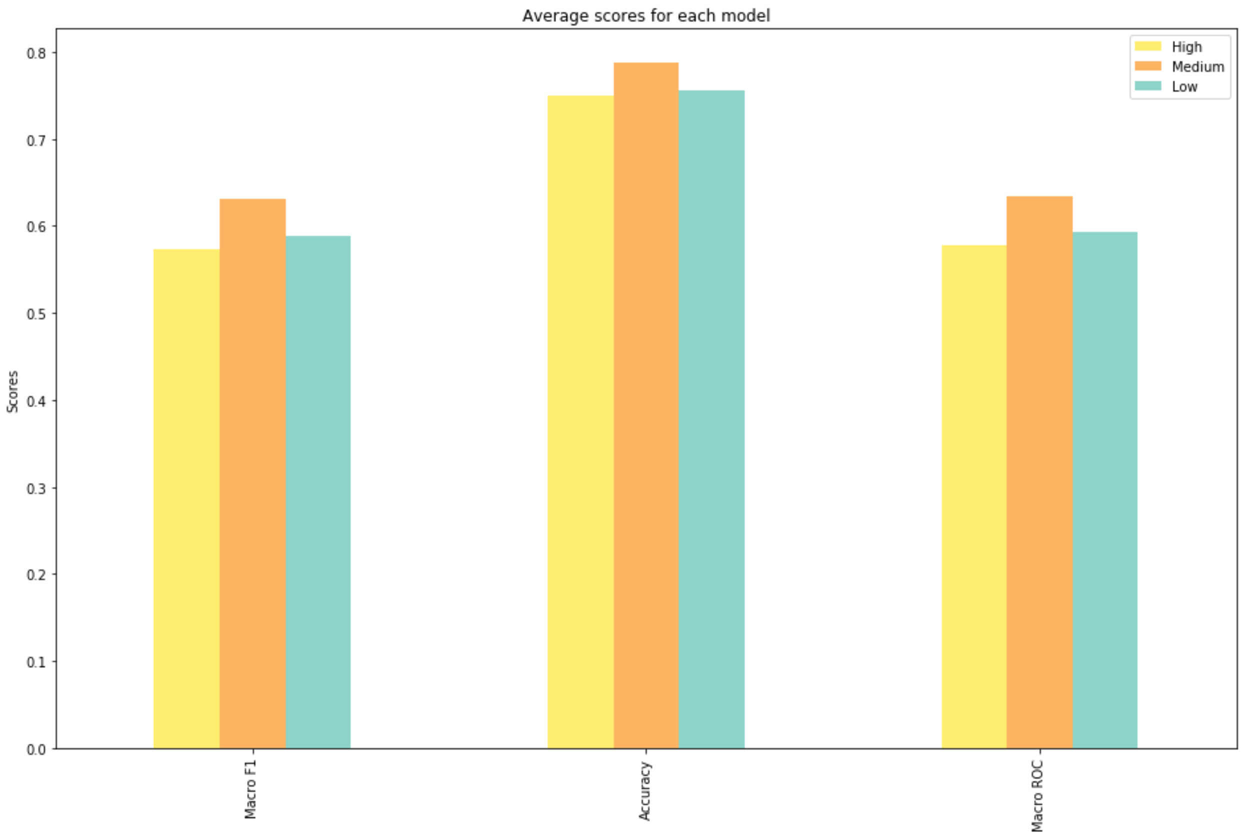


**Figure S2** (e): A comparison of performance of the classification results after binarization of the ANN output using different architectures.

Figure S2(e) shows the results after binarization of the ANN output. This also indicates that the ‘Medium’ architecture with three hidden layers is seen to outperform other architectures by all measures of performance. Hereafter, when we refer to ‘ANN’, we mean the ‘Medium’ architecture.

#### S2.2. Activation Functions

The output of ANN is determined using activation functions that can alter the outcome of the ANN drastically (Amari et al., 1997; Nwankpa et al., 2018). We conducted an experiment to compare various types of activation functions: Sigmoid, Softmax, Tanh, ReLu and Elu. These score comparisons when these functions were used are shown in Figure S2(f) and the results of the same are represented in Figure S2(g). We note from these graphs that ReLu outperformed the other activation functions and hence, Relu was used in our study.

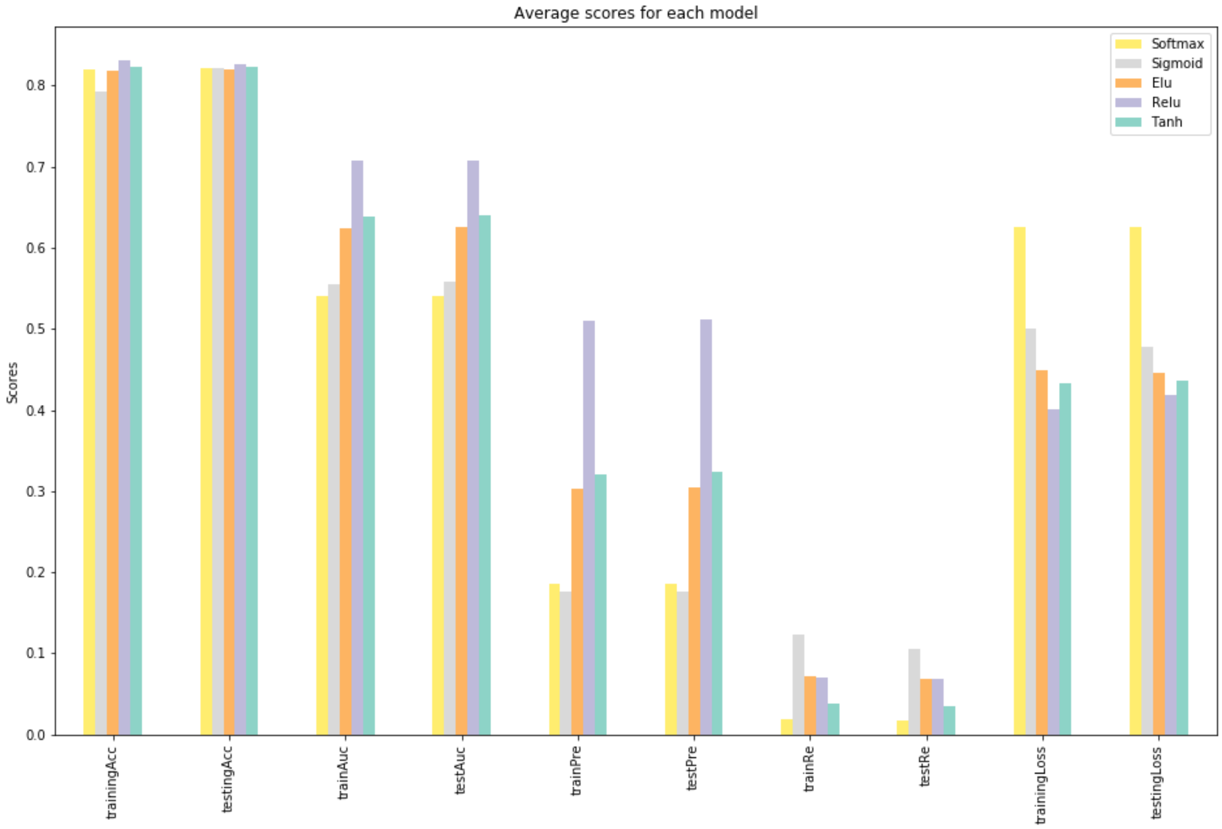


**Figure S2** (f): A comparison of the performance of the ANN with different activation functions.

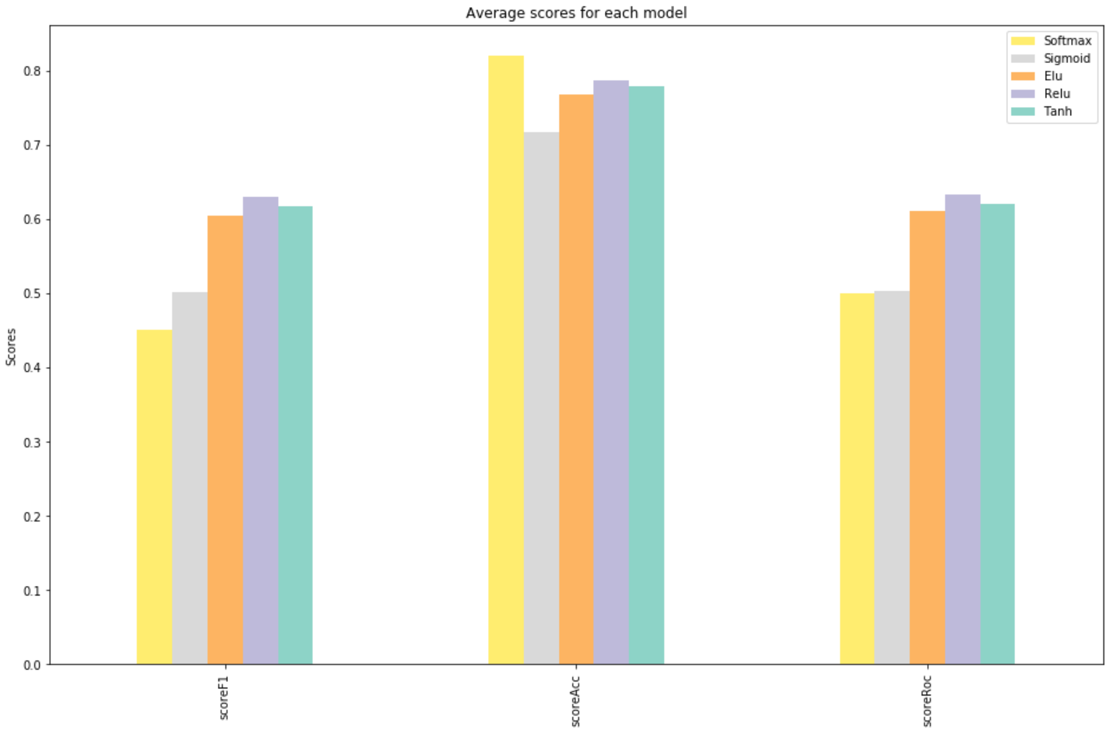


**Figure S2** (g): A comparison of the classification results after binarization of the ANN output using different activation functions.

**S2.3. Batch Size**

Batch size is one of the most important parameters to tune in modern deep learning systems. Practitioners often want to use a larger batch size to train their model as it allows computational speedups from the parallelism of GPUs. However, it is well known that too large a batch size will lead to poor generalization (G. Zhang et al., 2019). We experimented with seven batch sizes: 100, 500, 1000, 5000, 10000, 20000 and 30000. A comparison of the performance of the corresponding ANN models is presented in Figure S2(h).

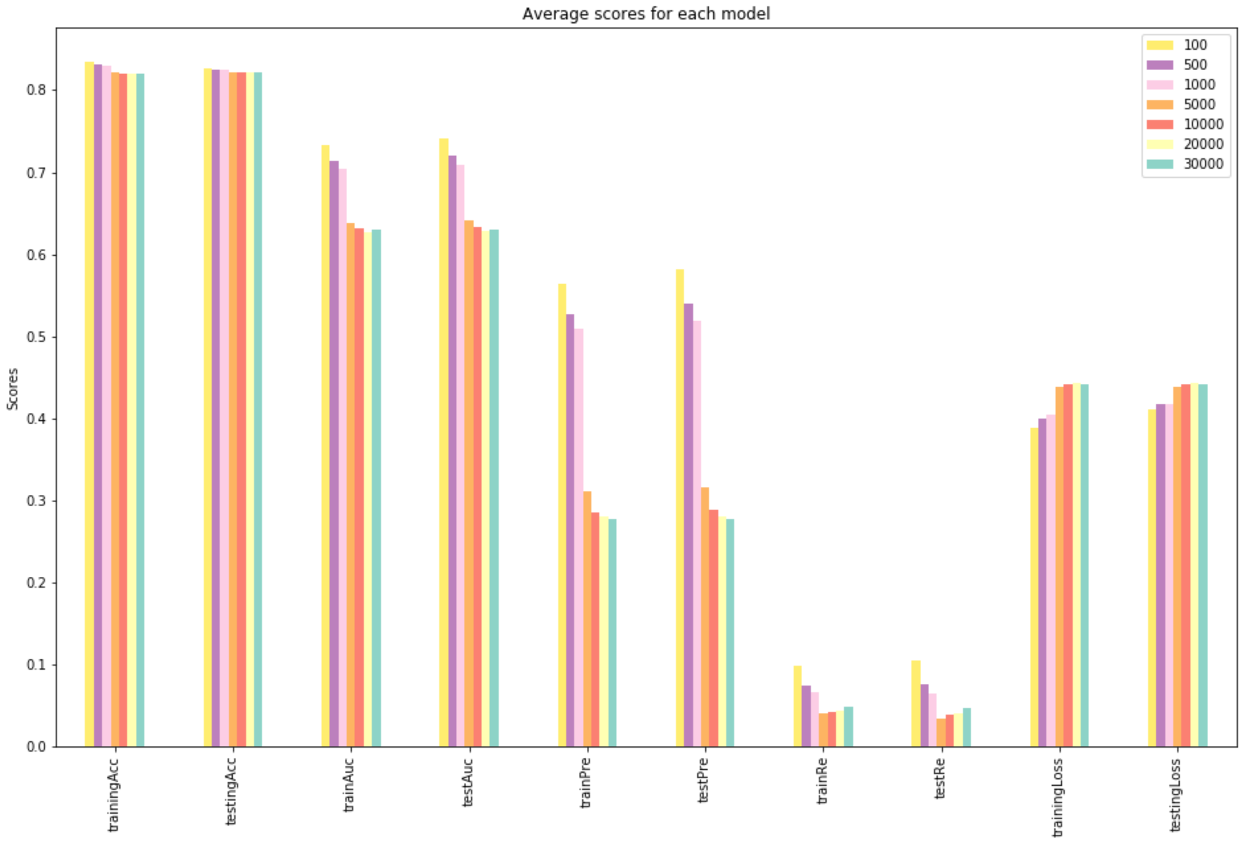


**Figure S2** (h): A comparison of the performance of ANN with  different batch sizes.

From this graph, we can see that there is a general decrease in trend with increase in the batch size. Hence, we can infer that smaller batch sizes give better overall results. To balance the compute time required due to the increased training time, we chose a batch size of 5000.

##### S2.4. Patience

The first sign of plateauing in model evaluation metrics is not the best time to stop training. This is because the model may get slightly worse or may coast into a plateau of no-further-improvement, before performing better (Amari et al., 1997). The experiment for selecting patience level was repeated 8 times with values 2, 3, 5, 8, 10, 13, 15 and 18. The performance of the corresponding ANN models using these patience levels are presented in Figure S2(i). It is evident from the results that the higher the patience, the better the results. A general increase in performance is observed with an increase in the patience level, but since higher patience translates to longer training times, we found 15 to be a good tradeoff.

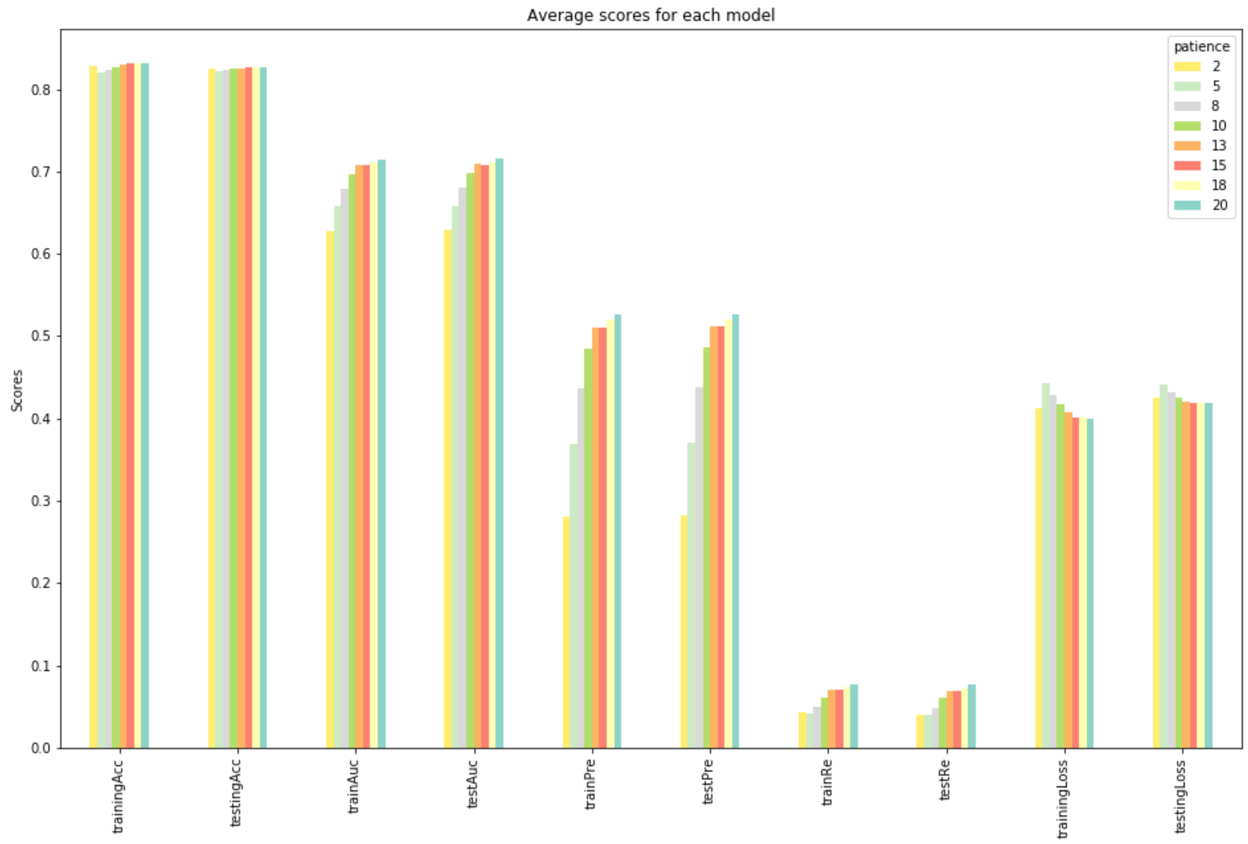


**Figure S2** (i): A comparison of the performance of ANN with different patience levels.

##### S2.5 The 10% Threshold for Data Point Selection

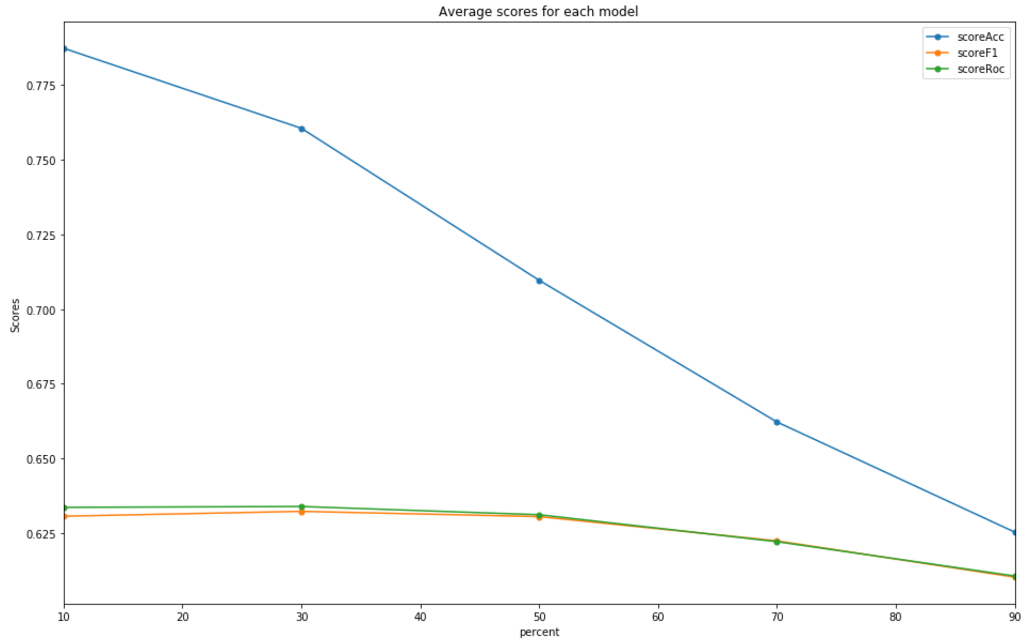


**Figure S2(j):** A comparison of the performance of ANN for multiple subsets, *viz.*, 10%, 30%, 50%, 70% and 90% of the data presenting the ADRs

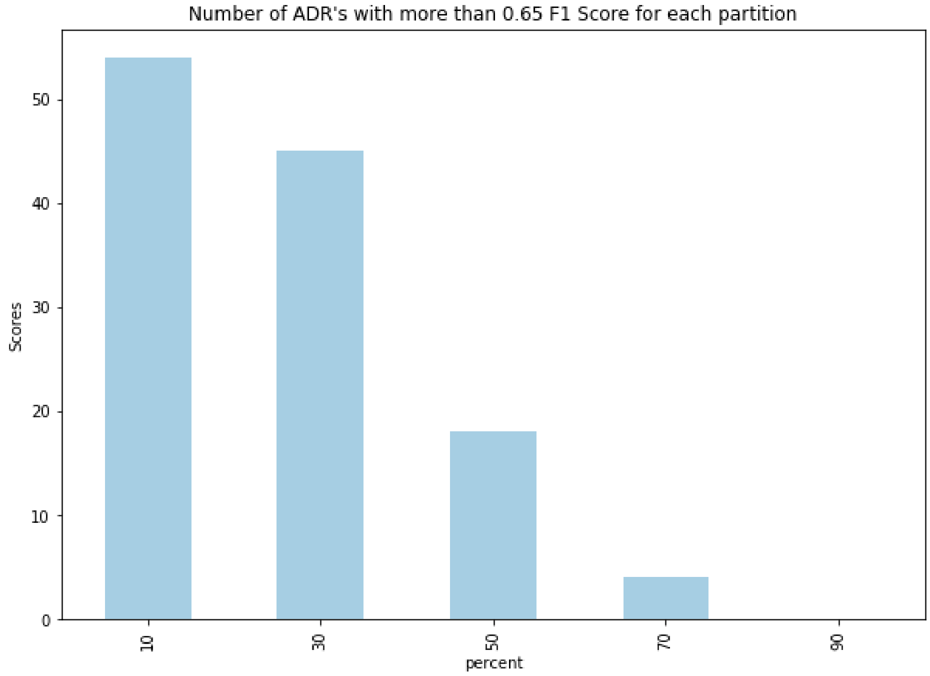


**Figure 4:** A comparison of the number of ADRs that can be predicted with an F1-score of greater than 0.65 for each partition of the data based on the percentage of data that presents the selected ADRs

**S3. Comparison of Various Machine Learning Models with ANNs**

A common pre-process to implementing many machine learning (ML) models mentioned below, is the dimensionality reduction using Principal Component Analysis (PCA) .(*Principal Component Analysis | I.T. Jolliffe | Springer*, n.d.) The input vector has 11,164 components. Many ML models are given to overfitting when there are a large number of features, especially when the number of samples are not significantly higher. To reduce the dimensions of the input PCA was used. PCA is based on a coordinate transformation of the input space to a space that best models the cumulative variance of the data. This is done through computing eigenvectors on the covariance matrix of the data. Each eigenvector is a linear combination of all the 11,164 components of the original input space. The weight of the component determines its importance for an eigenvector. Principal components are the eigenvectors that directions that best describe the variance in the data. The data is perfectly reconstructed when all 11,164 eigenvectors are used. However, PCA decorrelates the data and performs energy compaction that allows us to select a subset of the components. The subset of eigenvectors is selected on the basis of their importance---determined by the fraction of variance explained or the fractional contribution of the eigenvalue corresponding to the eigenvectors, selecting the eigenvectors corresponding to the largest of the eigenvalues. Through this subsampling, we achieve approximations of the data. We seek to achieve as close an approximation of the original data as possible, with as few components as possible through PCA. In our study, 1000 components were selected for model building as they represented 99% of the cumulative variance of the data.

With the input of 1000 features and using the concept of binary-relevance (M.-L. Zhang et al., 2018), the problem of multi-label prediction of ADRs for drug-pair combinations was reformulated into binary classification problems. We explored multiple classification approaches to solving these - namely KNN, Naive Bayes, Logistic Regression, extraTree (Geurts et al., 2006) and Artificial Neural Network. Figure S3(a) shows the performance of the various machine learning algorithms on the reduced dataset with 1000 Principal Components for 243 ADRs (the “10% dataset” -- see Section 2.1). From the plot, we note KNN, Logistic Regression and ExtraTree classifiers yielded the highest accuracies. However, their ROC was comparatively less, and hence, did not provide a good discriminative model. Naive Bayes, on the other hand, yielded a higher RoC but relatively lower Accuracy. Artificial Neural Network (ANN) provides promising results for all four evaluation metrics, even without significant effort tuning the model.

Figure S3(b) compares the result of ANN for 243 ADRs for a reduced dataset with 1000 Principal components against the entire dataset comprising 11,164 components for a patience level of 4. It is seen that the F1 Score and Precision, for the PCA based model, were higher than the original data without PCA by 3% and 4% respectively. This could be an indication that the PCA-based model is failing to label some of the ADRs. ANN without PCA performs better than on the reduced dimension space (after PCA), in terms Accuracy and ROC. It was evident that there are patterns of discriminatory value in the original dataset that may be lost in the reduced feature space. Further, considering that the ANN performs implicit feature selection, and that we may be able to achieve better performances with heuristic based tuning of ANN parameters on the training set, we opted to work with the original data without PCA.

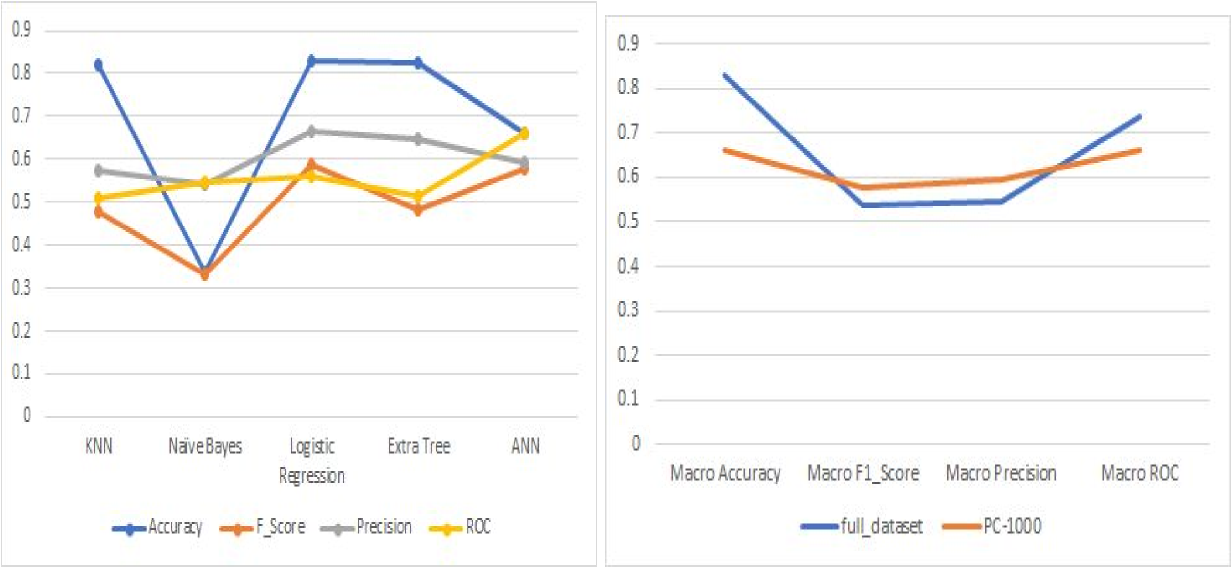


(a) (b)

**Figure S3** (a) The comparison of performance of five machine learning models for predicting 243 ADRs with 1000 principal components. (b) A comparison of the performance of ANN for the original input data with 11,164 components with that of an ANN with 1000 principal components (post PCA).

**S4. Evaluation Metrics for the ANN Model**

Macro averages of the three performance measures, viz., F1 score, Accuracy and RoC, across four validation sets and 243 ADRs is presented as a boxplot in Figure S4. Accuracy appears to be higher than the F1-score and RoC, as it accounts only for the positive predictions, not penalizing the system for false positives. F1 score (and, equivalently, the RoC) is a combination of both recall and precision that take into account false negatives and false positives respectively. Thus, these measures are lower. It is also the reason we chose a threshold that optimized the F1 score for each ADR on the training data, rather than accuracy.

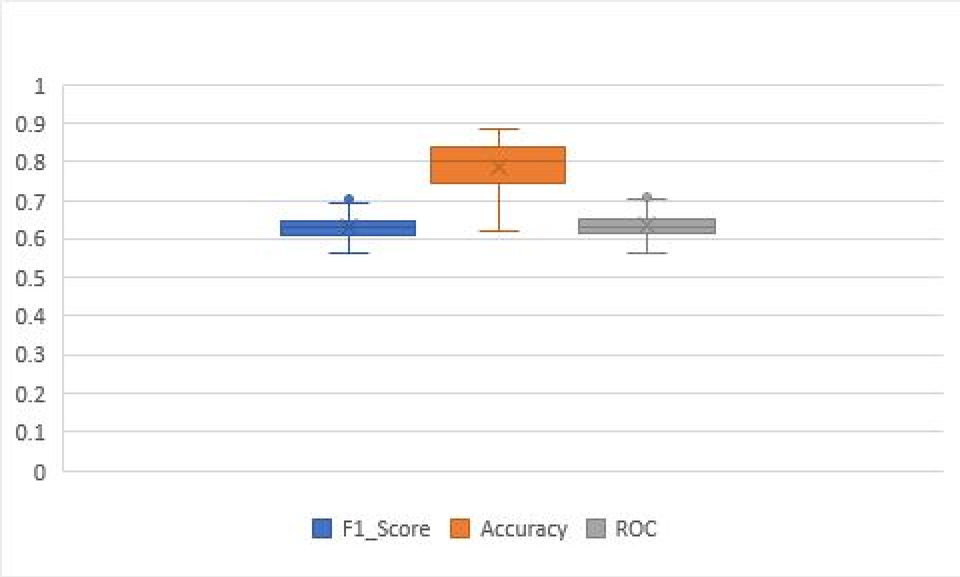


**Figure S4:** Boxplot showing the performance, based on Accuracy, F Score, Precision and ROC, for predicting 243 ADRs using ANN. The average of the performance metrics are: Accuracy - 0.7874; F Score - 0. 630536; ROC - 0.636511

**Table S1:** Minimum and maximum of each of the evaluation metrics out of 243 ADRs.

| Evaluation metrics | Maximum | ADR | Minimum | ADR |
|---|---|---|---|---|
| Accuracy | 0.885625 | Bursitis | 0.620875 | Difficulty Breathing |
| F_Score | 0.7039590 | Drug Addiction | 0.5617526 | Feeling Unwell |
| ROC | 0.7094963 | Bruxism | 0.5611701 | Feeling Unwell |
| **Maximum**: Highest evaluation metrics<br>**Minimum**: Lowest evaluation metrics | | | | |

**S5. Figshare details**

The files in Figshare that are required to run the code, and the respective output files are given below. The respective DOIs for each of the folders are also shared alongwith. Description of the dataset, and usage notes can be found in the README  for the respective scripts in the BitBucket repository.

2.1 Drug-pair Representation and the Associated ADRs (from the main text Section 2.1) has

- URL:https://figshare.com/articles/2_1_Drug-pair_Representation_and_the_Associated_ADRs/12579704
- DOI: 10.6084/m9.figshare.12579704

and contains:

1. ADR_list.csv
2. ADR_combine_hot_encoded.csv
3. ADR_combined.csv
4. Representation_of_single_drug.csv
5. required_ADR.csv
6. dataset_all.tar.gz containing:
    a. subset_1.tar.gz with
        i. ADR_dataset_for_training_subset_1.csv
        ii. ADR_validation_for_validation_subset_1.csv
        iii. dataset_subset_1.csv
        iv. dataset_validation_1.csv
    b. subset_2.tar.gz with
        i. ADR_dataset_for_training_subset_2.csv
        ii. ADR_validation_for_validation_subset_2.csv
        iii. dataset_subset_2.csv
        iv. dataset_validation_2.csv
    c. subset_3.tar.gz with
        i. ADR_dataset_for_training_subset_3.csv
        ii. ADR_validation_for_validation_subset_3.csv
        iii. dataset_subset_3.csv
        iv. dataset_validation_3.csv
    d. subset_4.tar.gz with
        i. ADR_dataset_for_training_subset_4.csv
        ii. ADR_validation_for_validation_subset_4.csv
        iii. dataset_subset_4.csv
        iv. dataset_validation_4.csv
    e. Main_dataset.tar.gz with
        i. ADR_combined.csv
        ii. dataset_subset.csv

- 3.2 Performance of the ANN Mode (from the main-text Section 3.2) has
    - URL: https://figshare.com/articles/3_2_Performance_of_the_ANN_Model/12582818
    - DOI: 10.6084/m9.figshare.12582818

and contains

1. Drug_pair_output_all_subsets.xlsx
2. S4. Evaluation Metrics for the ANN Model.csv

- Supplementary S3. Comparison of Various Machine Learning Models with ANNs-KNN_and_Naive_Bayes (from Supplementary material Supplementary section S3) has
    - URL:https://figshare.com/articles/Supplementary_S3_Comparison_of_Various_Machine_Learning_Models_with_ANNs-_KNN_and_Naive_Bayes/12582032
    - DOI: 10.6084/m9.figshare.12582032

and contains:

1. KNN_and_Naive_Bayes.tar.gz with
    a. KNN.tar.gz
    b. Naive_Bayes.tar.gz

- Supplementary S3. Comparison of Various Machine Learning Models with ANNs:extraTree_and_Logistic_Regression.tar.gz ( (from Supplementary material Section S3) has
    - https://figshare.com/articles/Supplementary_S3_Comparison_of_Various_Machine_Learning_Models_with_ANNs_extraTree_and_Logistic_Regression_tar_gz/12581228
    - DOI: 10.6084/m9.figshare.12581228

and contains:

a. extraTree_and_Logistic_Regression.tar.gz with
    i. extraTree.tar.gz
    ii. Logistic_Regression.tar.gz
